# Hippocampal Place Cells with NMDARs Do Not Require Excitation and Inhibition to Be Reciprocally Tuned

**DOI:** 10.64898/2026.04.27.721108

**Authors:** Samuel Gritz, Aaron D. Milstein

**Affiliations:** Center for Advanced Biotechnology and Medicine and Department of Neuroscience and Cell Biology, Rutgers Health, Rutgers, The State University of New Jersey, Piscataway, NJ 08854

## Abstract

In mouse hippocampal area CA1, excitatory pyramidal neurons referred to as “place cells” fire at specific locations in spatial environments during navigation. Many of the excitatory inputs to place cells are themselves spatially tuned, and prior work has shown that synaptic plasticity at those inputs strongly contributes to a selective increase in excitatory synaptic conductance when an animal is inside a cell’s “place field.” It is less clear whether inhibitory inputs to place cells vary with spatial position. Recent studies have investigated whether place cells receive spatially tuned inhibitory conductances by recording place cell activity *in vivo* and using computational models to help interpret experimental perturbations. One prior study used inhibitory optogenetics to suppress inhibitory neuron firing rates and observed a uniform depolarization of place cells across spatial locations, supporting a model with spatially uniform synaptic inhibition. In apparent conflict, other studies used excitatory optogenetics to depolarize place cells and observed a selective increase in excitability within place fields, supporting a model with a spatially localized decrease in inhibition. However, the latter studies overlooked the contribution of voltage-gated NMDA-type glutamate receptors (NMDARs) to synaptic integration, which are expected to contribute to the balance of excitatory and inhibitory synaptic currents. Here we show that when NMDARs are included at excitatory synapses in simple CA1 place cell models, all experimentally-observed properties of place cells can be recapitulated regardless of whether inhibition increases, decreases, or remains constant inside a place field.

**Significance Statement:** The hippocampus is a brain region required for the formation of new spatial and episodic memories (what happened where and when). Investigating the cellular and circuit mechanisms of memory recall could identify targets for therapies to combat memory decline associated with aging or neurodegeneration. Here we compare the results of computational models of the hippocampus to experimental recordings from mice to better understand the contribution of inhibitory neurons to the expression of spatial memories. We find that a special type of glutamate receptor, the NMDA receptor, helps to maintain the spatial selectivity of excitatory neurons in the hippocampus by counter-balancing fluctuations in the magnitude of inhibitory synaptic currents.

## Introduction

During spatial learning, excitatory neurons in the hippocampus called “place cells” develop location-specific firing patterns called “place fields,” which are thought to provide a cellular substrate for spatial and episodic memory (O’Keefe and Conway, 1978; Nakazawa et al., 2004; Buzsaki and Moser, 2013). A primary contributor to the spatial selectivity of place cells is an increase in the strengths of excitatory synaptic inputs that are active inside a cell’s place field, which are shaped during learning by synaptic plasticity (Nakazawa et al., 2004; Bittner et al., 2017; Milstein et al., 2021; Li et al., 2024; Gonzalez et al., 2025). It is less clear whether the amount of inhibitory synaptic input to a place cell is spatially tuned, or whether such modulation plays an important role in determining the properties of place cells (Grienberger et al., 2017a; Weber and Sprekeler, 2018; Geiller et al., 2020; Geiller et al., 2022; Jeong and Singer, 2022; Valero et al., 2022b; Valero et al., 2022a). For a place cell to receive a specific increase or decrease in inhibitory synaptic conductance within its place field would require that it selectively connect to interneurons that either increase or decrease their firing rates at those spatial positions. However, compared to excitatory neurons, inhibitory interneurons in the hippocampus have higher baseline firing rates and are less spatially tuned, with a fraction of interneurons expressing either moderate increases or decreases in activity at some spatial positions (McNaughton et al., 1983; Marshall et al., 2002; Grienberger et al., 2017a; Geiller et al., 2020; Hainmueller et al., 2024).

Interestingly, regardless of whether inhibitory synaptic conductance varies across spatial locations, when a place cell depolarizes inside its place field, the electrical driving force on chloride ions through GABA(A) receptors (GABARs) increases, which is expected to increase the magnitude of inhibitory synaptic currents selectively within the place field (Grienberger et al., 2017a). In a recent study, Valero et al. (Valero et al., 2022a) investigated the spatial tuning of inhibition using a combination of experimental perturbations and computational model simulations. They reasoned that if inhibitory synaptic currents were larger inside place fields, then the response of place cells to extrinsic depolarizing stimulation would be reduced in-field relative to out-of-field. Counter to this prediction, they observed that artificial depolarization of place cells with optogenetic stimulation actually resulted in an increased response within place fields (Valero et al., 2022a) (Fig. 1C). A simple single-compartment computational model of a place cell receiving excitatory AMPA-type glutamate receptor (AMPAR) and inhibitory GABAR conductances suggested that this experimentally-observed increase in excitability inside place fields could only occur if inhibitory conductance selectively decreased within the place field (Fig. 1D), but not if it increased or remained constant.

**Figure 1.**
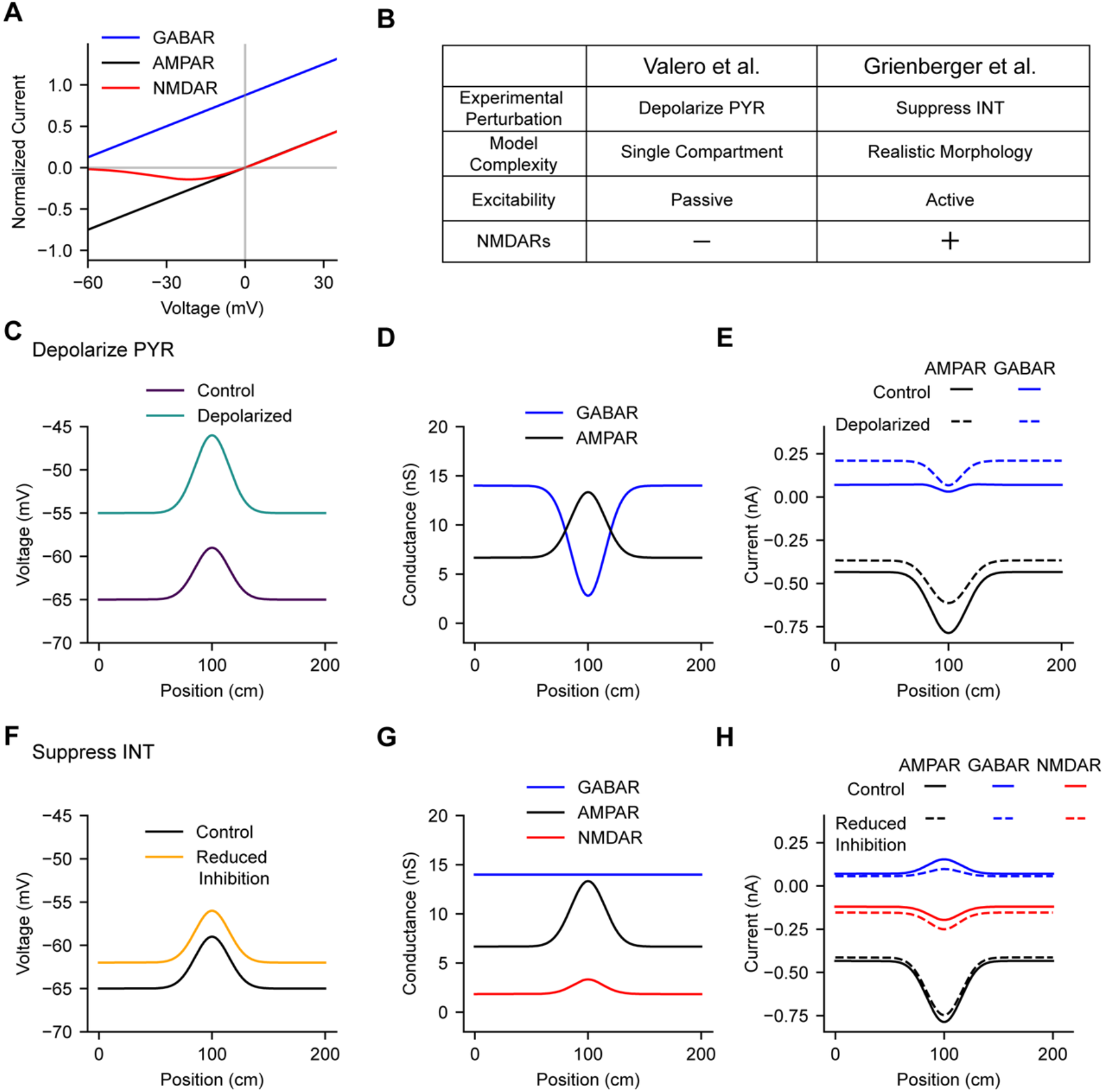
**A**, Current-voltage relationships are shown for AMPARs, NMDARs, and GABARs. Voltage-dependent NMDARs increase current at depolarized voltages, whereas AMPAR currents decrease and GABAR currents increase due to changes in ionic driving force. **B**, Table summarizes experiments and models reported in Valero et al. (Valero et al., 2022a) and Grienberger et al. (Grienberger et al., 2017b). **C**, In Valero et al., excitatory pyramidal place cells were artificially depolarized by optogenetic stimulation, revealing higher excitability in-field than out-of-field. **D**, The experimental results in C were recapitulated by a simple single-compartment place cell model without NMDARs and with GABAR conductance (blue) that decreased within the place field. **E**, Synaptic currents measured from the model in D are shown before (solid lines) and after (dashed lines) artificial depolarization. Artificial depolarization decreases AMPAR currents (black) and causes larger increases in GABAR current out-of-field compared to in-field (blue). **F**, In Grienberger et al., the firing rates of inhibitory interneurons were suppressed optogenetically, causing uniform depolarization of place cells in-field and out-of-field. **G**, The experimental results in F were recapitulated by a multi-compartment biophysical model with NMDARs (red) and with spatially uniform GABAR conductance (blue). **H**, Synaptic currents measured from the model in G are shown before (solid lines) and after (dashed lines) suppression of inhibition. In this model, GABAR and NMDAR currents increase in-field, while AMPAR currents decrease.

In apparent conflict with the above result, Grienberger et al. (Grienberger et al., 2017a) investigated the spatial tuning of inhibition with a different experimental perturbation – optogenetic suppression of CA1 inhibitory interneuron firing – and found that reducing synaptic inhibition onto place cells resulted in a uniform depolarization of place cells both in-field and out-of-field (Fig. 1F). A morphologically- and biophysically-detailed computational model of a place cell suggested that this experimental observation was consistent with a constant level of spatially untuned inhibition arriving at all spatial locations (Fig. 1G). In that model, both AMPARs and voltage-dependent NMDA-type glutamate receptors (NMDARs) were included at excitatory synapses, and their conductances were calibrated to reproduce experimental results showing that NMDARs contribute to the nonlinear summation of excitatory synaptic inputs in pyramidal neurons (Losonczy and Magee, 2006; Harnett et al., 2012; Grienberger et al., 2017a). Thus, increases in inhibitory GABAR currents in-field resulting from changes in chloride driving force were counterbalanced by increases in excitatory NMDAR currents as the cell depolarized (Fig. 1H). NMDARs are known to be expressed in CA1 place cells (Racca et al., 2000), where they have been shown to contribute to burst firing (Grienberger et al., 2014), network synchrony and population oscillations (Buzsaki, 2002; Leung and Shen, 2004; Hunt and Kasicki, 2013), place field formation (Bittner et al., 2017; Grienberger and Magee, 2022), and memory formation (Nakazawa et al., 2004).

In the study by Valero et al. (Valero et al., 2022a), models with NMDARs were not tested, and the models without NMDARs were not validated against the experimental perturbation data from Grienberger et al. (Grienberger et al., 2017a) where synaptic inhibition was suppressed. On the other hand, the study by Grienberger et al. (Grienberger et al., 2017a) did not test models with alternative spatial patterns of synaptic inhibition, and did not validate the uniform inhibition model against the experimental perturbation data from Valero et al. (Valero et al., 2022a) where place cell excitability was probed with extrinsic depolarization. Here we systematically tested simple CA1 place cell models with and without NMDARs receiving inhibitory conductances that were either spatially uniform, balanced, or reciprocally tuned, and we optimized and validated the models against all available experimental perturbation data. We found that when voltage-gated NMDARs are included at excitatory synapses, they counterbalance changes in inhibitory driving force and enable all known properties of CA1 place fields to be recapitulated regardless of the spatial distribution of synaptic inhibition.

## Materials and Methods

### CA1 Place Cell Model

Place cell membrane potential dynamics in response to spatially- and temporally-modulated synaptic inputs during simulated treadmill running (Fig. 2) were modeled using custom Python code, which is accessible on GitHub (Gritz and Milstein). The membrane voltage of a single-compartment neuron model evolved over time according to:

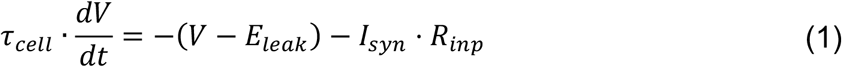

where *τ*_*cell*_ = 20 *ms* represents a membrane time constant, *R*_*inp*_ = 100 *M*Ω corresponds to input resistance, and *E*_*leak*_ is the reversal potential of a leak conductance calibrated to enforce an out-of-field membrane potential ~−63 mV. The total synaptic current received by the CA1 place cell, *I*_*syn*_, was modeled as the sum of excitatory (AMPAR and NMDAR) and inhibitory (GABAR) synaptic currents:

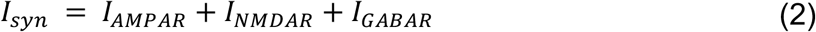

**Figure 2.**
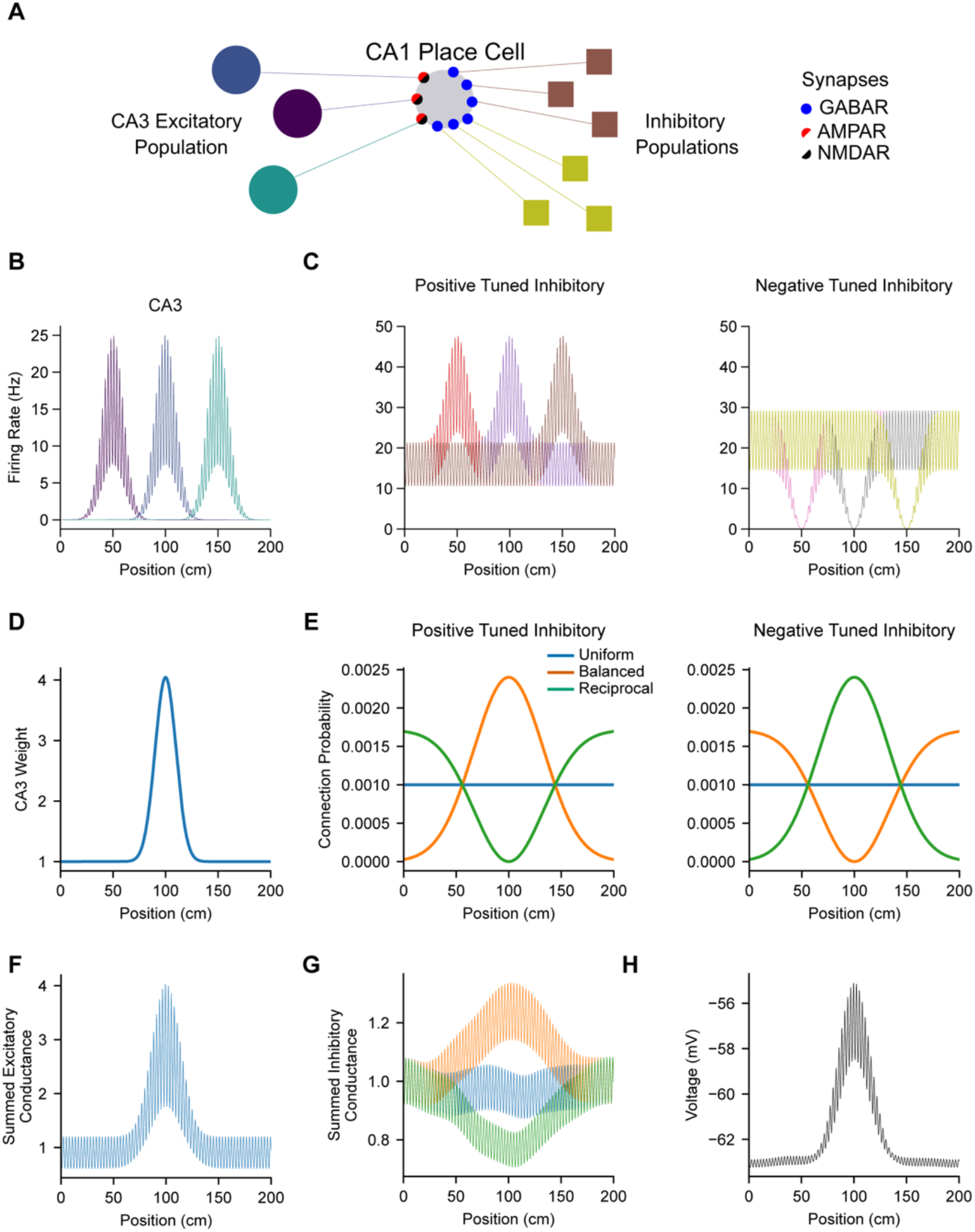
**A**, Diagram depicts synaptic inputs to a CA1 place cell. In all models, excitatory inputs from CA3 place cells contact synapses containing AMPARs (black), and inhibitory inputs from two subpopulations of CA1 interneurons contact synapses containing GABARs (blue). In a subset of models, excitatory synapses receiving CA3 inputs also contain NMDARs (red). **B**, The firing rates of CA3 place cell inputs to the model CA1 place cell were theta-modulated and spatially-tuned. **C**, The firing rates of CA1 inhibitory interneurons were theta-modulated and spatially-tuned. Two subpopulations of interneurons were either positively (left) or negatively (right) modulated by spatial position. **D**, Spatially-tuned depolarization of the CA1 place cell model was achieved by modulating the synaptic strength of AMPAR synapses from CA3 place cells depending on their place field locations. **E**, Three possible configurations of synaptic inhibition (uniform, blue; balanced, orange; reciprocal, green) were compared by varying the connection probabilities of the two subpopulations of spatially-tuned interneurons to the model CA1 place cell. **F**, The weighted sum of CA3 excitatory inputs (B and D) generates a spatially- and temporally-modulated excitatory AMPAR conductance. **G**, The sum of CA1 interneuron inhibitory inputs (C and E) generates a spatially- and temporally-modulated inhibitory GABAR conductance. **H**, Time-varying excitatory and inhibitory synaptic currents resulting from the conductances in F and G are integrated by the model CA1 cell’s membrane, resulting in a spatially- and temporally-modulated intracellular membrane voltage (V_m_). Traces shown in B, C, and F-H reflect a representative single trial from a model with uniform inhibition and without NMDARs.

Note that, per convention, excitatory inward (depolarizing) currents have a negative sign. At a given type of synapse *X* = {*AMPAR, NMDAR, GABAR*}, synaptic current *I*_*X*_ was calculated as:

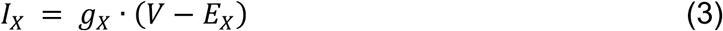

where *g*_*X*_ is the total synaptic conductance and *E*_*X*_ is the reversal potential for that type of synaptic ion channel (*E*_*AMPAR*_ = 0 *mV, E*_*NMDAR*_ = 0 *mV, E*_*GABAR*_ = −70 *mV*). Voltage-dependent modulation of synaptic currents through NMDAR ion channels by Mg^2+^ ions was approximated by a sigmoidal voltage-dependent factor (Spruston et al., 1995):

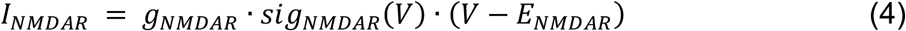

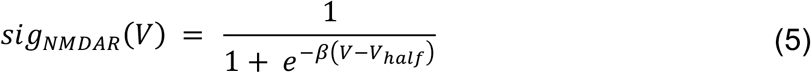

where *β* determines the slope and *V*_*half*_ determines the threshold (Supp. Table S1).

The temporal dynamics of synaptic conductances *g*_*X*_ depended on a weighted sum of presynaptic firing rates and an intrinsic decay with a time constant specific to each synapse type (*τ*_*AMPAR*_ = 5 *ms, τ*_*NMDAR*_ = 75 *ms*, and *τ*_*GABAR*_ = 5 *ms*):

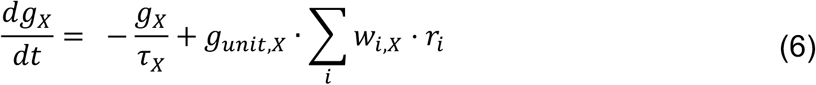

where *g*_*unit,X*_ is a unitary conductance scaling factor (Supp. Table S1), *w*_*i,X*_ is the relative strength of the connection from presynaptic neuron *i*, and *r*_*i*_ is the firing rate of presynaptic neuron *i*. To produce a place field in the model CA1 cell (Fig. 2H), excitatory AMPAR weights from a population of CA3 place cell inputs were modulated as a Gaussian function of spatial position with a floor width of 60 cm (Fig. 2D), a minimum value of 1 and a maximum value of 1 + Δ*W*_*AMPAR*_ optimized for each model (Supp. Table S1). All excitatory NMDAR synaptic weights were set to 1 for models with NMDARs, and were set to 0 for models without NMDARs. All inhibitory GABAR synaptic weights were set to 1.

### Spatial Modulation of Presynaptic Inputs

A population of 1000 CA3 place cells provided excitatory input to the model CA1 place cell. Each CA3 input expressed a Gaussian-shaped place field with a floor width of 60 cm, a minimum out-of-field firing rate of 0 Hz, and a maximum in-field firing rate of 25 Hz. The preferred locations of each cell in the population were spaced equally across a 200 cm circular track (Fig. 2B). Two subpopulations of local CA1 interneurons provided inhibitory inputs to the model CA1 place cell, one with positive spatial tuning, and one with negative spatial tuning. Spatial modulation for both types of inhibitory neurons was Gaussian shaped with a floor width of 60 cm, with either peak or trough locations spaced equally across the track. The firing rates of the two populations were calibrated to each have a mean rate of 25 Hz, resulting in positively-tuned interneurons ranging from a minimum rate of 21.3 Hz out-of-field to a maximum rate of 47.7 Hz at the peak of their spatial modulation, and negatively-tuned interneurons ranging from a maximum rate of 29.2 Hz out-of-field to a minimum rate of 0 Hz at the trough of their spatial modulation (Fig. 2C).

For each model variant, the shape of the summed inhibitory conductance to the CA1 place cell model (uniform, balanced, or reciprocal) was determined by modulating the probability of connecting with interneurons in the two inhibitory subpopulations (Fig. 2E and 2G). Spatially uniform inhibition was achieved by equally and uniformly sampling connections from both interneuron subtypes (Fig. 2E, blue). Balanced inhibition was achieved by preferentially connecting to positively-tuned interneurons with peak locations close to the place field location of the CA1 place cell, and preferentially connecting to negatively-tuned interneurons with trough locations far from the CA1 cell’s place field (Fig. 2E, orange). Reciprocal inhibition was achieved in the opposite manner, by preferentially connecting to positively-tuned interneurons with peaks far from the target cell’s field, and by preferentially connecting to negatively-tuned interneurons with troughs nearby (Fig. 2E, green). Connectivity preferences were implemented as Gaussian functions of position with a floor width equal to the full track length (200 cm). For each model variant, the total number of connections made from each interneuron population were calibrated such that the average summed inhibitory conductances from the two subpopulations were approximately equal (Supp. Table S1). Each model variant was evaluated by simulating and averaging across five instances of the model with different instantiations of the random connectivity.

### Temporal Modulation of Presynaptic Inputs

The firing rates of all presynaptic populations were modulated by a 7 Hz theta rhythm. For each trial type (with or without simulated experimental perturbations), simulations were conducted for five trials, and on each trial, a global reference oscillation analogous to an extracellularly-recorded local field potential (LFP) was offset by a random phase *ϕ*_*offset*_ to decouple theta phase from spatial position. Each presynaptic population was assigned a theta modulation depth *d*, and a preferred phase of firing *ϕ*_*pop*_ (Supp. Table S1). Then, their spatial firing rates were multiplied by a temporal modulation factor *M*_*pop*_. For the inhibitory neuron populations (Fig. 2C), *M*_*INH*_ was defined as:

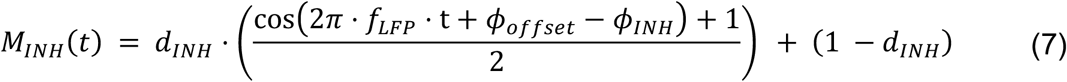

where *f*_*LFP*_ = 7 *Hz*, and *d*_*INH*_ = 0.5. For the excitatory CA3 population, the frequency of each neuron’s theta modulation and its phase relative to the LFP-like reference varied with spatial position. This spatial relationship between the phase of firing and spatial position is a form of temporal coding referred to as phase precession (O’Keefe and Recce, 1993; Skaggs et al., 1996; Harvey et al., 2009; Mizuseki et al., 2009; Geisler et al., 2010; Bittner et al., 2015; Chadwick et al., 2015; Grienberger et al., 2017a). CA3 place cells were modeled as phase precessing a full 360° during place field traversal, with their phase relative to the LFP crossing *ϕ*_*CA*3_ in the center of their fields (Supp. Table S1). This was achieved using a previously reported method (Chadwick et al., 2015), summarized here:

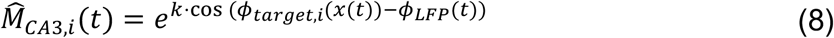

where *k* = 0.7 is a factor that determines the shape of the firing rate modulation by theta phase, *x*(*t*) is the spatial position along the track, *ϕ*_*target,i*_(*x*) is the target firing phase at each spatial position for presynaptic CA3 neuron *i*, and *ϕ*_*LEP*_(*t*) is the phase of the LFP reference. Finally, temporal modulation *M*_*CA*3,*i*_ for CA3 place cell *i* is computed as:

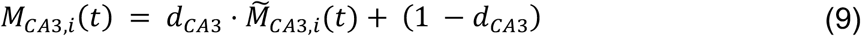

where 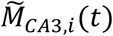 reflects a normalization of 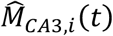 to span the range from 0 to 1, and *d*_*CA*3_ = 0.7. As previously reported (Geisler et al., 2010; Grienberger et al., 2017a), this results in the postsynaptic CA1 place cell expressing an intracellular voltage oscillation that is close to *f*_*LEP*_ out of field and increases in magnitude and frequency in field, resulting in phase precession of the peaks of the intracellular voltage oscillation relative to the LFP reference, consistent with experimental observations (Harvey et al., 2009; Grienberger et al., 2017a).

### Mimicking Experimental Perturbations

In Valero et al. (Valero et al., 2022a), place cells were experimentally depolarized by optogenetic stimulation. To mimic this perturbation, we ran a series of simulations with varying amounts of extrinsic current applied to the model place cell, calibrated to drive out-of-field membrane voltage to a range of target values (Fig. 3F). In Grienberger et al. (Grienberger et al., 2017a), synaptic inhibition onto place cells was experimentally reduced by optogenetically suppressing the activity of CA1 inhibitory interneurons. This perturbation was simulated by reducing all synaptic inhibitory conductances by 50%.

**Figure 3.**
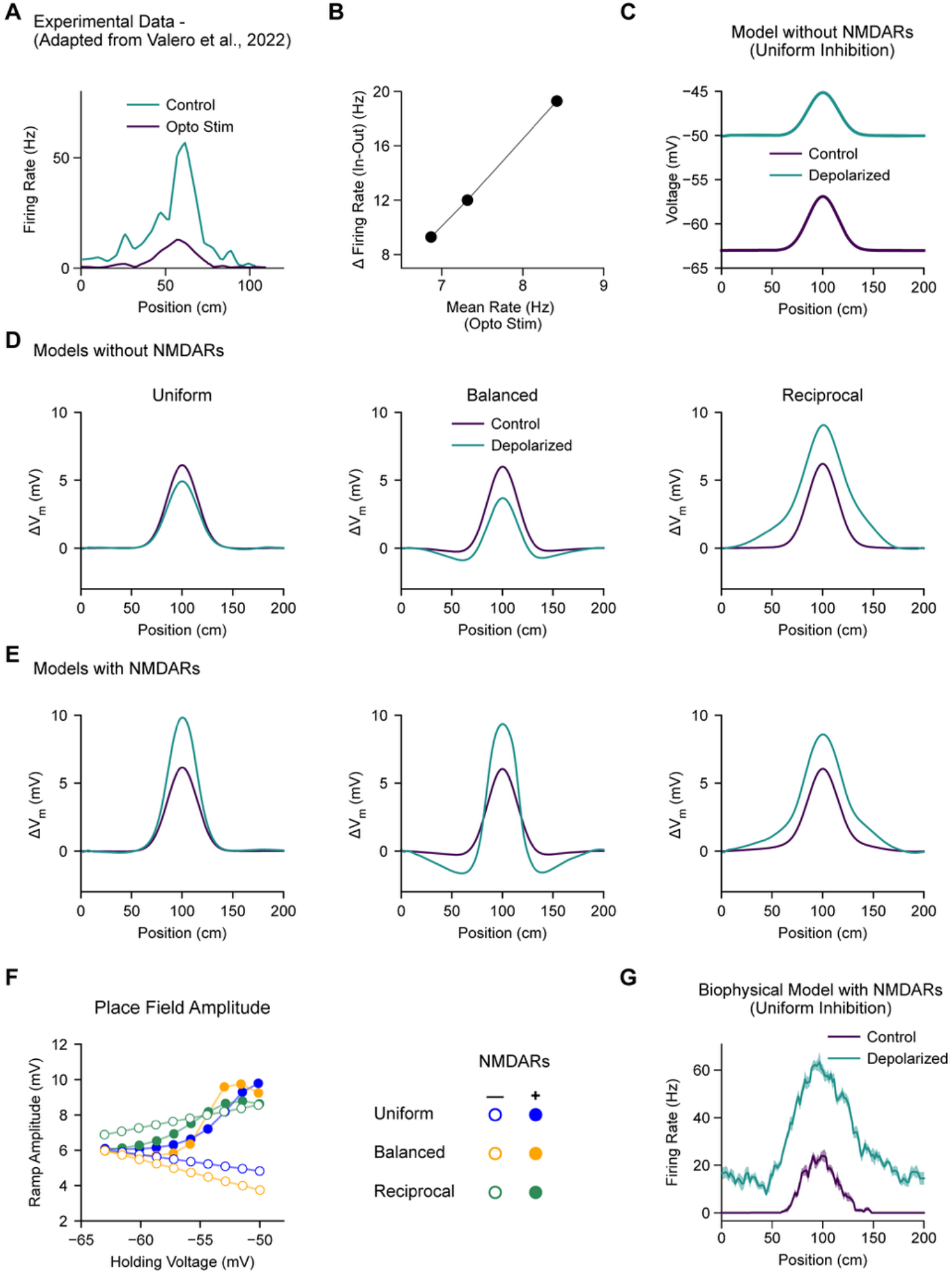
**A-B**, Experimental data adapted from Valero et al. (Valero et al., 2022a). **A**, Average firing rate of an example place cell with (teal) and without (purple) artificial depolarization by optogenetic stimulation. **B**, Across a range of stimulation amplitudes, the response to stimulation was larger in-field than out-of-field. **C**, Simulating the experimental perturbation from A in a simple CA1 place cell model with uniform synaptic inhibition and without NMDARs does not recapitulate the experimental result. **D**, The relative change in V_m_ relative to out-of-field is shown for place cell models without NMDARs with three different configurations of inhibition: uniform (left), balanced (middle), and reciprocal (right). Only the reciprocal model recapitulates the experimental result in A. **E**, Same as D for models with NMDARs. All three models recapitulate the experimental result in A. In C-E, shading reflects SEM across trials. **F**, The amplitude of intracellular place field ramp depolarization is quantified across a range of holding voltages for the six models in D and E. Error bars reflect standard deviation across five independent instances of each model. **G**, The biophysical place cell model with uniform inhibition and with NMDARs reported in Grienberger et al. (Grienberger et al., 2017b) also recapitulates the experimental result in A. Shading reflects SEM across trials.

### Measuring Place Field Properties

Properties of model place cell intracellular membrane potential (V_m_) dynamics were measured by first filtering voltage time series data from single trial simulations with a low-pass filter (<2 Hz) to analyze the slow subthreshold ramp depolarization that underlies CA1 place fields, and with a bandpass filter (4 – 10 Hz) to analyze properties of the intracellular theta oscillation (Bittner et al., 2015; Bittner et al., 2017; Grienberger et al., 2017a). Theta oscillation envelope time series were generated using the scipy implementation of the Hilbert transformation (Oppenheim and Schafer, 2010; Virtanen et al., 2020). Filtered voltage ramp and theta envelope time series were then trial-averaged, and measurements of in-field and out-of-field depolarization and theta amplitude were made by averaging values within 0.4 s windows. To quantify phase precession of the intracellular theta oscillation relative to the LFP reference, peaks were detected from the theta bandpass-filtered traces of individual trials, and then a trial-averaged map of theta phase versus spatial position was used to compute the extent of phase precession within a 4 s window surrounding the peak of the place field. For each model variant, the values of all measured features were then averaged across five network instances with different random connectivity.

### Model Parameter Optimization

Six model variants (uniform, balanced, and reciprocal inhibition, each with or without NMDARs) were optimized by using a population-based iterative multi-objective search within bounds specified for 9 free parameters (Deb and Jain, 2013; Milstein, 2021) (Supp. Fig. S1 and Supp. Table S1). Objectives were calculated for 30,000 variants of each model based on distance of the following model features to experimentally-derived target values: the V_m_ outside the place field (target: −63 mV), the amplitude of the place field subthreshold ramp depolarization (target: 6 mV), the mean phase of intracellular theta oscillation peaks outside the field (target: 180°), the difference in the magnitude of intracellular theta oscillations in-field versus out-of-field (target: >0.5 mV), the extent of intracellular theta phase precession (target: >100°), the effect of experimentally perturbating place cell V_m_ on place field ramp amplitude (Valero et al., 2022a) (target slope: >0.23 mV/mV), and the effects of experimentally perturbing synaptic inhibition on place cell V_m_ and intracellular theta oscillations (Grienberger et al., 2017a), including the change in V_m_ inside and outside the field (target: >1.5 mV), the mean phase of intracellular theta oscillations outside the field (target: 180°), and the increase in intracellular theta envelope amplitude inside and outside the field (target: >0.2 mV). The change in the extent of phase precession induced by perturbing synaptic inhibition was not explicitly optimized, but was included as a held-out feature for evaluating models against an experimental target of <-25%. Each objective was computed as the squared error relative to its target value, normalized by a specified tolerance term, and averaged across five instantiations of each network with different random connectivity. Most objectives were treated as one-sided “soft” objectives, penalized only when the value measured from the model undershot the target, with the exception of out-of-field V_m_, place field ramp amplitude, and the mean theta phase before and after inhibitory perturbation. Following optimization, in addition to the “best” models with parameters yielding the lowest total objective error, we also analyzed degenerate models with parameters that were large distances from the “best” model, but yielded similarly low objective errors. In Table 1 and Supp. Fig. S2, we evaluated each model variant based on feature values that were averaged across five members of these families of degenerate models with alternative parameters (a.k.a. “Marder groups” (Milstein et al., 2022)). This demonstrates that the results we report for place cell models with different spatial modulation of synaptic inhibition with and without NMDARs reflect general features of those model configurations and do not depend critically on the specific choice of hyperparameters.

**Table 1.**
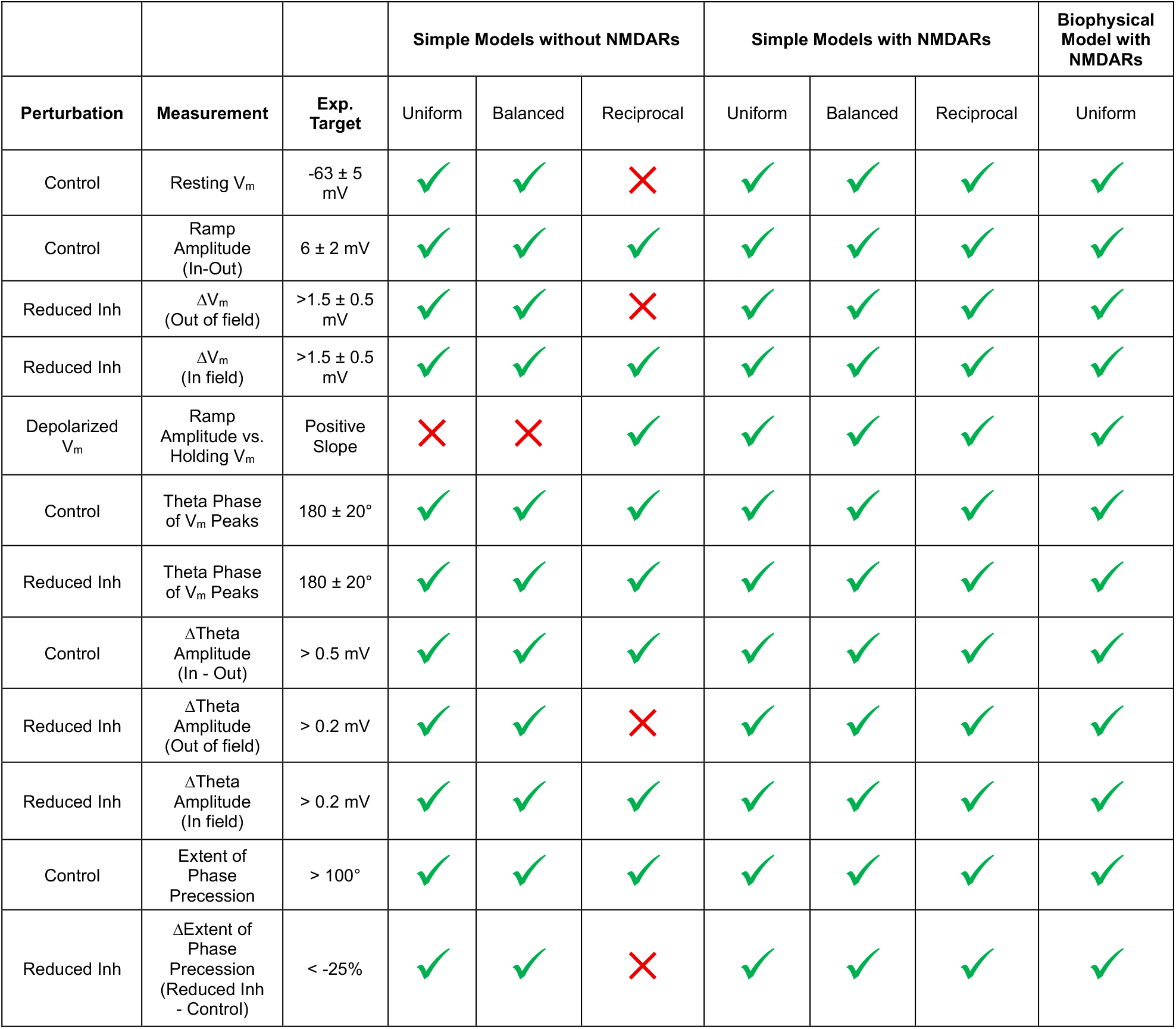
Each CA1 place cell model configuration was evaluated against criterion based on experimental targets. For each model configuration, five alternative models with different parameters were evaluated (see Methods). Green checkmarks indicate ≥ 4/5 model variants met criterion. Red crosses indicate ≤ 1/5 model variants met criterion.

## Results

We first constructed a simple, single-compartment model of the intracellular voltage dynamics of a CA1 place cell receiving rate-based synaptic inputs from excitatory CA3 place cells and two subpopulations of local GABAergic inhibitory interneurons, one with moderate positive spatial tuning, and one with moderate negative spatial tuning (Fig. 2A-C, Methods). To achieve a spatially localized increase in membrane voltage (V_m_) at a pre-determined place field location in the CA1 place cell model (Fig. 2H), the synaptic weights of CA3 inputs were set as a Gaussian function of the distance between their presynaptic place field locations and the place field location of the postsynaptic CA1 place cell (Fig. 2D, Methods), as has been observed experimentally *in vivo* (Bittner et al., 2017; Gonzalez et al., 2025). To achieve synaptic inhibitory conductances that either increased, decreased, or remained constant within the CA1 cell’s place field, we varied the connection probability with inhibitory interneurons based on the locations of their positive or negative spatial modulation (Fig. 2E, Methods). This resulted in model variants with either uniform (blue), balanced (orange), or reciprocal (green) spatial modulation of total inhibitory conductance (Fig. 2G). Inputs to the model were also temporally modulated by a 7 Hz theta oscillation (Buzsaki, 2002) (Fig. 2B and 2C). This enabled phasic interactions between excitatory and inhibitory synaptic currents to produce an intracellular V_m_ oscillation (Fig. 2H), which we compared to experimental data from *in vivo* intracellular recordings of place cells (Harvey et al., 2009; Bittner et al., 2015; Grienberger et al., 2017a; Valero et al., 2022b). In particular, place cell recordings have shown that intracellular theta oscillation amplitude, frequency, and phase are modulated by spatial position (Harvey et al., 2009; Bittner et al., 2015) and are sensitive to perturbation of synaptic inhibition (Grienberger et al., 2017a; Valero et al., 2022b). Each model variant was optimized to match a variety of target features derived from experimental recordings of place cell V_m_ dynamics (Supp. Fig. S1 and S2, Methods).

We then took these three optimized model variants with different spatial distributions of inhibition and challenged them with a depolarizing stimulus mimicking the optogenetic activation of place cells performed in Valero et al (Valero et al., 2022a) (Fig. 3A-C). Examining models that expressed only AMPARs at excitatory synapses (and not NMDARs), we corroborated the previous findings that only the model with reciprocal inhibition (Fig. 3D, right), but not models with uniform (Fig. 3D, left) or balanced inhibition (Fig. 3D, middle), recapitulated the experimentally observed increase in place field amplitude, which scaled with increasing background depolarization (Fig. 3F, open circles). However, when NMDARs were added to excitatory synapses and the models were re-optimized to match known place field properties, all model variants (uniform, balanced, and reciprocal inhibition) exhibited increasing place field amplitude with increasing background depolarization (Fig. 3E and 3F), matching the experimental perturbation data (Fig. 3A and 3B). We also performed the same test on the morphologically- and biophysically-detailed model with uniform inhibition and NMDARs used in Grienberger et al. (Grienberger et al., 2017a) and found that model also recapitulated the experimentally-observed increase in excitability within the place field. These results demonstrate that when NMDARs are present at excitatory synapses, their voltage-dependent conductance can provide a source of inward current that compensates for the loss of AMPAR current (Losonczy and Magee, 2006; Poleg-Polsky and Diamond, 2016) and counterbalances the increase in GABAR current that occurs with depolarization within a CA1 cell’s place field. Thus, reciprocal inhibition is not a unique configuration, and it is not required to explain the results of the place cell V_m_ perturbations reported in Valero et al (Valero et al., 2022a).

Next, we sought to evaluate how these six model variants responded to a perturbation of synaptic inhibition similar to that performed experimentally by Grienberger et al (Grienberger et al., 2017a). In that study, the spatial waveform of depolarization in place cells as well as properties related to the intracellular theta oscillation were measured before and after optogenetically suppressing inhibitory interneurons (Fig. 4C). They reported that under control conditions, the magnitude of intracellular theta oscillations increased within the place field relative to outside the field (Fig. 4C, middle, black). Then, upon transient suppression of inhibition, the mean V_m_ and the intracellular theta amplitude increased both in-field and out-of-field (Fig. 4C, left and middle). These experimental observations established important features of place cell V_m_ dynamics that we used as targets for model optimization (Table 1) to ensure that the parameters controlling the relative magnitudes of excitatory and inhibitory synaptic conductances lay within a biologically plausible range (Methods).

**Figure 4.**
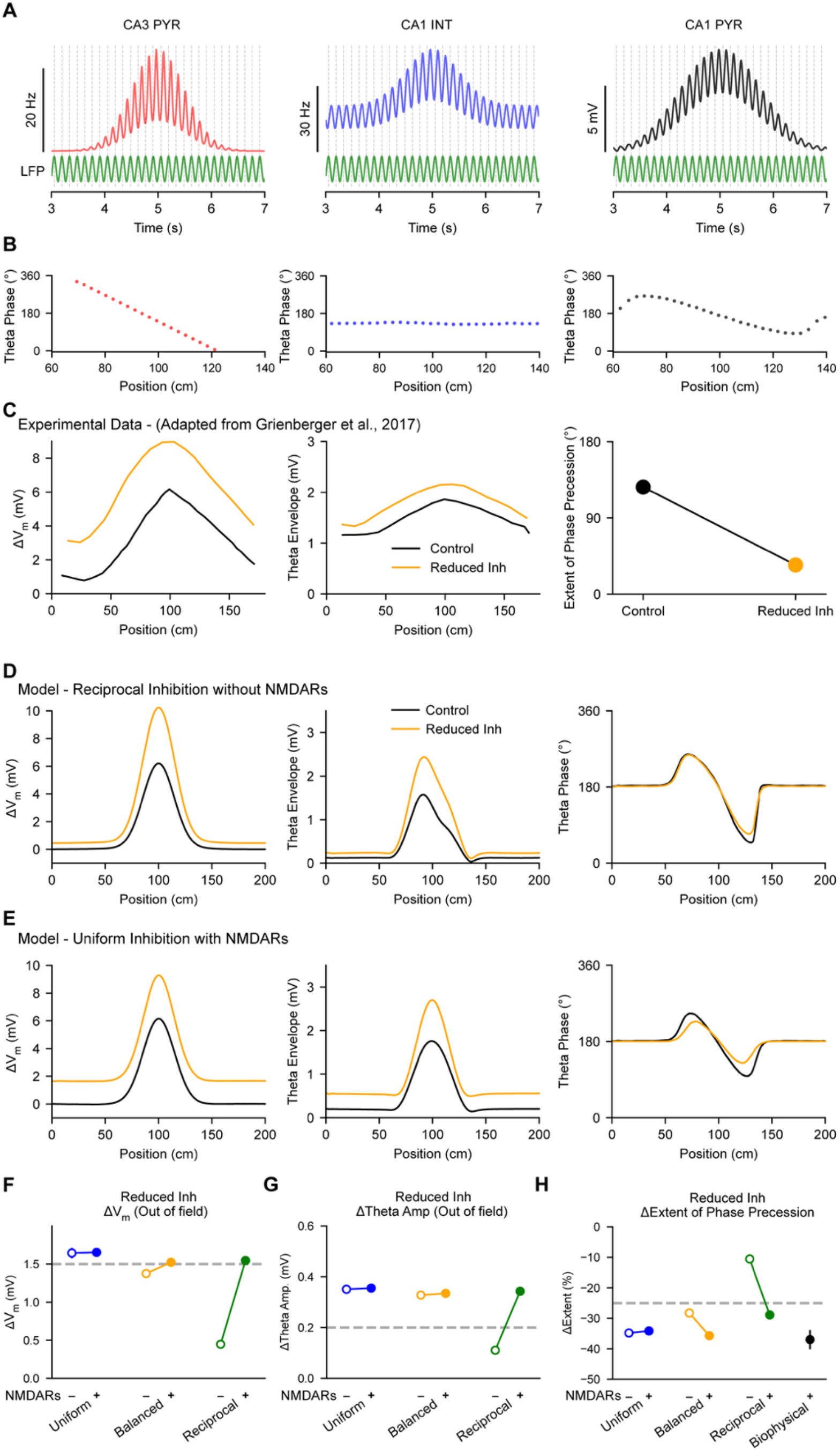
**A**, Theta-modulated firing rates are shown for examples of model excitatory CA3 place cell inputs (left) and inhibitory CA1 inputs (middle), and theta-modulated intracellular V_m_ is shown for the model CA1 place cell (right). **B**, The theta phases of firing rates or V_m_ peaks shown in A are calculated relative to an external LFP reference (green), and plotted versus spatial position. CA3 inputs phase precess through their place fields (left), while CA1 interneurons maintain a constant phase (middle). Interactions between these inputs result in phase precession in the CA1 place cell (right). **C**, Experimental data adapted from Grienberger et al. (Grienberger et al., 2017b). Left: The change in V_m_ (relative to resting V_m_) of a CA1 place cell recorded *in vivo* is shown before (black) and after (orange) optogenetic suppression of inhibition. Middle: same as left for the envelope of theta-filtered V_m_. Right: Same as left for the extent of phase precession of CA1 intracellular V_m_ peaks. **D**, Simulation results are shown from a CA1 place cell model with reciprocal inhibition and without NMDARs, for comparison to C. **E**, Same as D for a model with uniform inhibition model and with NMDARs. **F-H**, Effects of simulating optogenetic suppression are compared across six model configurations. Optimization targets are shown with a dashed line. **F**, The change in out-of-field theta amplitude is quantified. **G**, The change in out-of-field V_m_ is quantified. **H**, The change in the extent of phase precession of intracellular V_m_ peaks is quantified. Results from a biophysically detailed model with uniform inhibition and with NMDARs from Grienberger et al. is shown for comparison. In D and E, shading reflects SEM across trials. In F-H, error bars reflect standard deviation across five independent instances of each model.

Another interesting property of CA1 place cell V_m_ dynamics is that, relative to an extracellular local field potential (LFP) reference, the frequency of intracellular theta oscillations transiently increases within the place field. This results in the peaks of individual cycles of the intracellular V_m_ oscillation (as well as any resulting action potentials) occurring progressively earlier in theta phase as the animal moves through the yplace field (Fig. 4A and 4B, right). This phenomenon is referred to as “theta phase precession” and underlies a form of temporal coding that enables information about the spatial position of an animal to be encoded in both the rate and the timing of spikes emitted by a place cell (O’Keefe and Recce, 1993; Skaggs et al., 1996; Mizuseki et al., 2009). Here we modeled phase precession similarly to Grienberger et al. (Grienberger et al., 2017a) – as constructive interference between inhibitory currents oscillating at the same frequency as the extracellular LFP (Fig. 4A and 4B, middle), and excitatory currents resulting from a weighted sum of CA3 place cell inputs that were themselves phase precessing within their place fields (Geisler et al., 2010; Jaramillo et al., 2014; Chadwick et al., 2015; Grienberger et al., 2017a) (Fig. 4A, left). Interestingly, Grienberger et al. (Grienberger et al., 2017a) also observed that the extent of theta phase precession was markedly reduced upon perturbation of synaptic inhibition (Fig. 4C, right), and this feature was recapitulated by their biophysical model with uniform inhibition and NMDARs. We used this result as a held-out feature for evaluation of each of the six models after optimization (Table 1).

Analysis of the optimized models revealed that only the model with reciprocal inhibition without NMDARs failed to recapitulate a variety of experimentally measured features of place cell V_m_ dynamics (Fig. 4D and Table 1). In particular, that model did not meet target criterion for the following responses of place cells to perturbation of synaptic inhibition: out-of-field increase in V_m_ (Fig. 4D, left and Fig. 4F), out-of-field increase in intracellular theta amplitude (Fig. 4D, middle and Fig. 4G), and in-field decrease in the extent of phase precession (Fig. 4D, right and Fig. 4H). In contrast, all models with NMDARs, including the model with uniform inhibition (Fig. 4E), succeeded in matching all experimentally derived targets (Fig. 4F-H and Table 1). Interestingly, the uniform and balanced inhibition models without NMDARs, despite failing to reproduce the in-field excitability result from Valero et al. (Valero et al., 2022a) (Fig. 3), succeeded in replicating the inhibitory perturbation results from Grienberger et al (Grienberger et al., 2017a) (Fig. 4F-H, Table 1). Importantly, the properties of each model configuration that we report here were shared across families of models with alternative parameters identified during optimization (Supp. Fig. S1 and S2, Methods), indicating that these properties are general consequences of each model’s structure rather than resulting from particular choices of parameters. Overall, these results demonstrate generally that, as long as place cells express NMDARs, they do not require spatially tuned inhibitory conductance, and specifically that place cell models with reciprocal tuning of inhibition and without NMDARs are inconsistent with experimentally observed responses to perturbation of synaptic inhibition.

## Discussion

In this study, we systematically tested the responses of simple models of hippocampal CA1 place cells to simulated experimental perturbations to better understand the contributions of spatially tuned synaptic inhibition and voltage-dependent NMDARs to the properties of place fields. We found that including NMDARs at excitatory projections from CA3 to CA1 place cells conferred an insensitivity to the spatial tuning of inhibition such that a battery of experimentally measured features of place cell intracellular voltage dynamics could be recapitulated regardless of whether synaptic inhibition increased, decreased, or remained constant within place fields. While these results indicate that place cells do not require synaptic excitation and inhibition to be reciprocally tuned, they imply that the properties of place cells would be robust to differences in the spatial profile of synaptic inhibition that may occur due to heterogeneity in connectivity or changes that occur during learning (Geiller et al., 2020; Geiller et al., 2022; Jeong and Singer, 2022).

Interestingly, theoretical work has shown that when inhibitory neurons receive random recurrent connections from both excitatory and inhibitory neurons, moderate stimulus selectivity can emerge through network dynamics, and this can be sufficient to shape the stimulus selectivity of excitatory neurons without requiring biased or structured connectivity or inhibitory synaptic plasticity (Milstein et al., 2022; Galloni et al., 2026). Thus, uniform and random connectivity between excitatory and inhibitory neurons in the hippocampus remains the simplest model consistent with experimental observations that does not require that specific mechanisms be in place to achieve a precise wiring configuration. Interestingly, prior work combining functional imaging and anatomical tracing has indicated that, in the hours following the initial formation of a new place field through excitatory synaptic plasticity, the hippocampal network may reorganize to incorporate the new place cell into subnetworks of excitatory and inhibitory neurons with preferential wiring (Geiller et al., 2022). Structured connectivity has also been observed in cortical regions, where substantial experimental and theoretical evidence supports a model of balanced excitation and inhibition achieved through preferential connectivity with interneurons that share the stimulus tuning of their excitatory targets (Anderson et al., 2000; Isaacson and Scanziani, 2011).

The models presented in this study predict that currents through voltage-dependent NMDARs counterbalance increased GABAR currents when place cells are depolarized within their place fields, maintaining a high excitatory-inhibitory (EI) ratio and elevated excitability. Directly testing this prediction experimentally *in vivo* could be challenging, but may be possible with recently developed tools. First, an existing place cell must be identified for targeted intracellular recording, or a place field must be experimentally induced in a silent cell by evoking behavioral timescale synaptic plasticity (BTSP) (Bittner et al., 2017; Milstein et al., 2021; Rolotti et al., 2022), which requires that NMDARs be intact. Then, ideally place field V_m_ dynamics would be recorded both before and after NMDARs are blocked selectively in the recorded cell, which could be achieved using DART (drug acutely restricted by tethering) pharmacology (Shields et al., 2024). Such a manipulation combined with extrinsic depolarization may be able to discriminate between the multiple possible configurations of synaptic inhibition (Valero et al., 2022a).

A caveat of the modeling presented in this study is that the neuronal cell model does not express any voltage-gated ion channels other than NMDARs, and contains only one single compartment, which restricts all synapses to sensing the same limited membrane voltage range shared by the soma. In biological neurons with extensive dendritic arbors, spines, and active conductances, voltage compartmentalization is expected to add heterogeneity to the voltages sensed by different NMDARs at different synapses, and changes in driving force may be more or less pronounced depending on the synapse location (Harnett et al., 2012; Beaulieu-Laroche and Harnett, 2018; Lafourcade et al., 2022). Nonetheless, our results suggest that the subthreshold voltage dynamics of place cells are well approximated when nonlinear NMDAR activation is included.

Separate from the question of whether stimulus tuned inhibition contributes to receptive field properties is the related question of how synaptic inhibition contributes to the gating of excitatory plasticity in pyramidal neurons (Wilmes et al., 2016; Milstein et al., 2021; Jeong and Singer, 2022; Rolotti et al., 2022; Campbell et al., 2026). Some experiments have indicated that heterogeneities in circuit wiring that pre-exist before learning can bias when plasticity occurs and in which cells (McKenzie et al., 2021). Other theoretical work has suggested that there may be functional advantages to learning stimulus selectivity in the subtypes of inhibitory neurons that are responsible for gating plasticity in excitatory neurons (Galloni et al., 2026). Understanding whether or how specific interneuron cell types influence memory formation and recall will be critical to design interventions for memory deficits associated with disrupted EI balance, including epilepsy and Alzheimer’s disease (Okechukwu et al., 2026).

## Acknowledgments

We are grateful for early discussions with Jeff Magee and Sandhya Senthilkumar. Large-scale compute resources were provided by NSF-supported allotments from TACC (Frontera) and ACCESS (Bridges-2). We are grateful for funding provided by Rutgers Biomedical and Health Sciences and NIMH grant R01MH135576.

## Supplementary Information

**Supplementary Figure S1.**
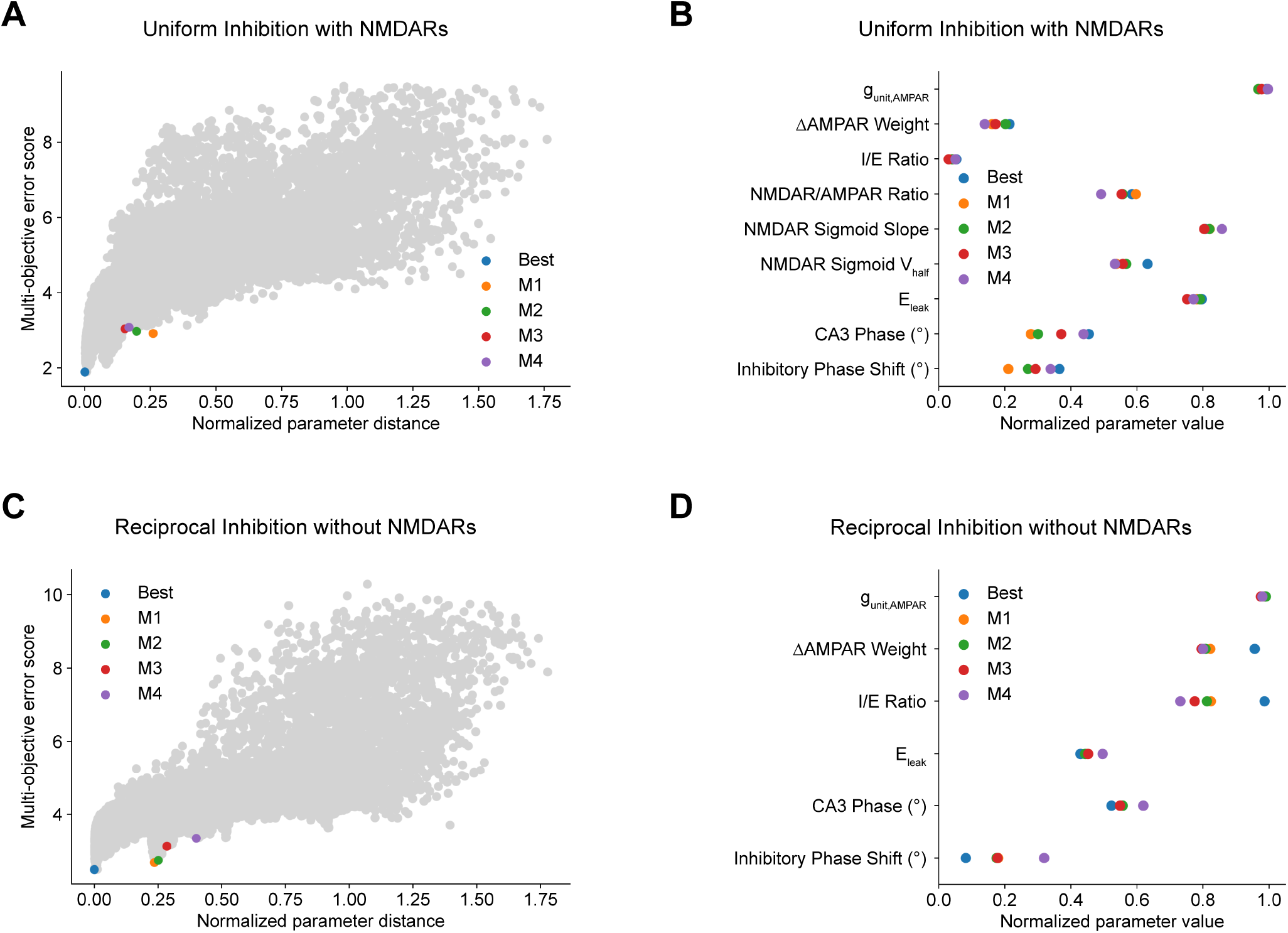
**A**, During parameter optimization, CA1 place cell model variants with different parameters were evaluated against criterion based on experimental targets. A family of models were chosen that performed similarly across optimization criterion, but exhibited diversity in their parameter values. Shown are model variants of the model configuration with uniform inhibition and with NMDARs. **B**, Parameter values are shown for the five model variants labeled in A. **C-D**, Same as A-B for the model configuration with reciprocal inhibition and without NMDARs.

**Supplementary Figure S2.**
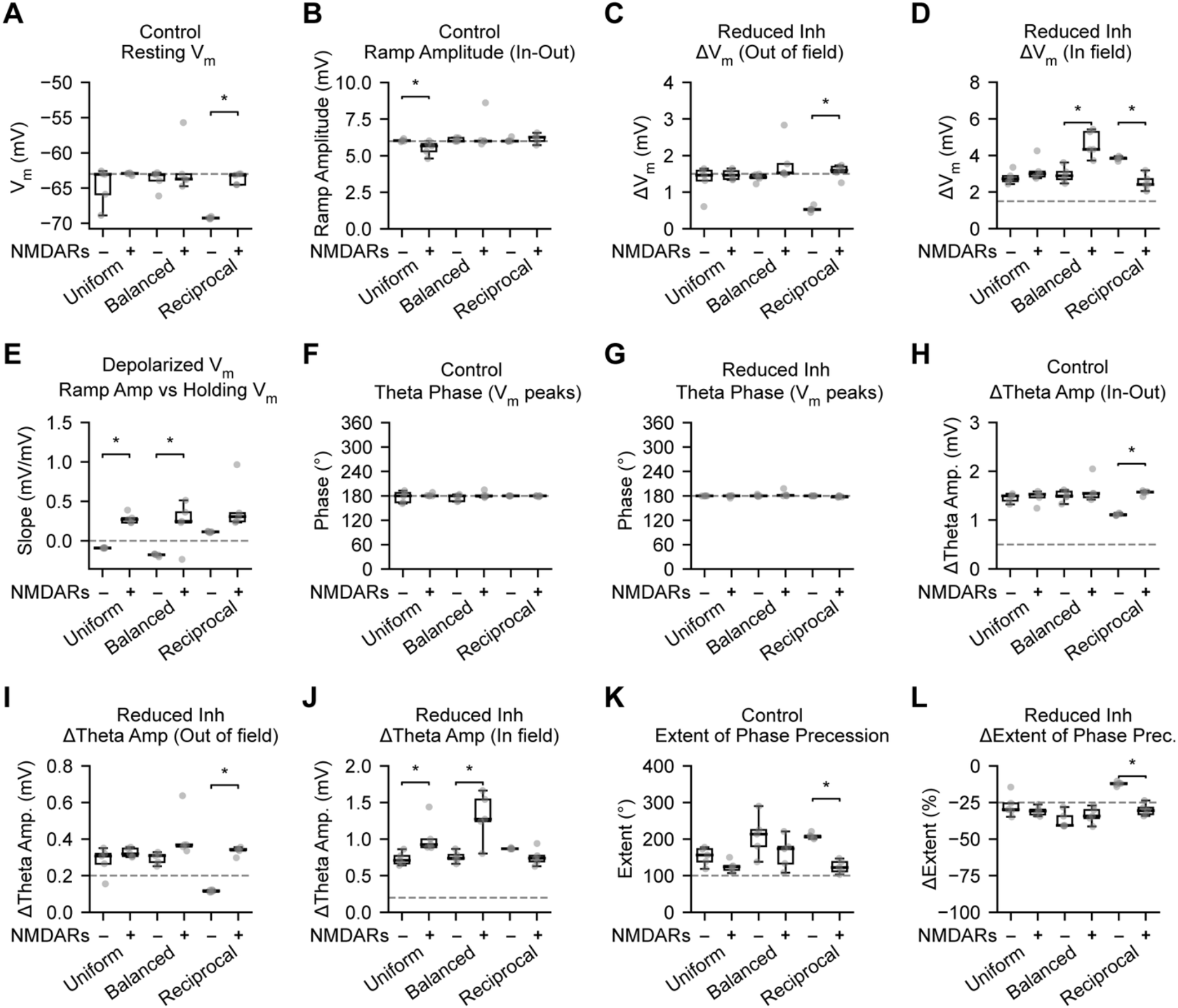
**A-L**, For each CA1 place cell model configuration, feature values were measured and compared against experimental targets for five variants of each model with different alternative parameters. Data is displayed as box-and-whisker plots with individual data points, group medians and inter-quartile ranges indicated.

**Supplementary Table S1.**
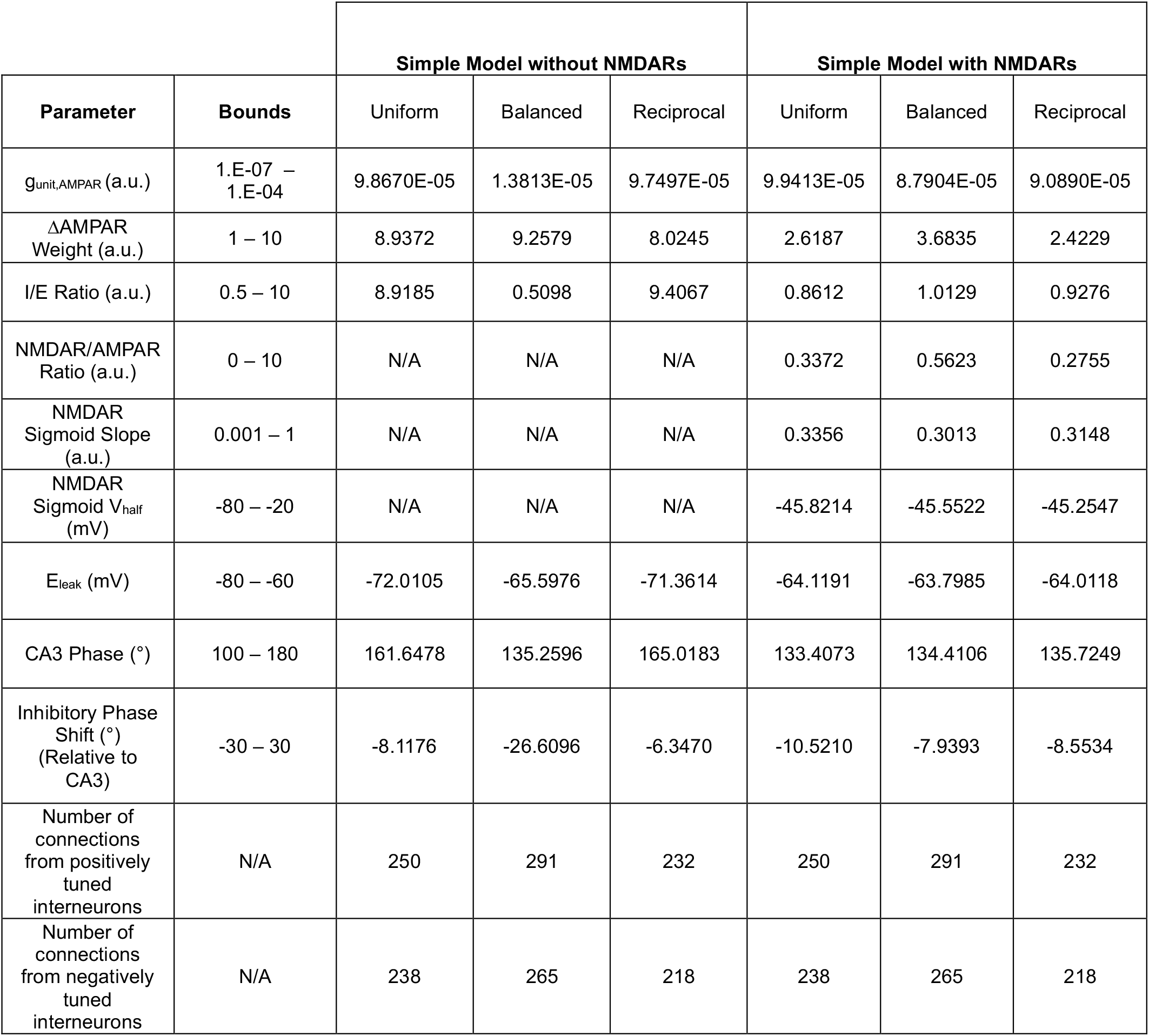
Bounds and optimized parameters are shown for six CA1 place cell model configurations.

## References

Anderson JS, Carandini M, Ferster D (2000) Orientation tuning of input conductance, excitation, and inhibition in cat primary visual cortex. J Neurophysiol 84:909–926.

Beaulieu-Laroche L, Harnett MT (2018) Dendritic Spines Prevent Synaptic Voltage Clamp. Neuron 97:75–82 e73.

Bittner KC, Milstein AD, Grienberger C, Romani S, Magee JC (2017) Behavioral time scale synaptic plasticity underlies CA1 place fields. Science 357:1033–1036.

Bittner KC, Grienberger C, Vaidya SP, Milstein AD, Macklin JJ, Suh J, Tonegawa S, Magee JC (2015) Conjunctive input processing drives feature selectivity in hippocampal CA1 neurons. Nat Neurosci 18:1133–1142.

Buzsaki G (2002) Theta oscillations in the hippocampus. Neuron 33:325–340.

Buzsaki G, Moser EI (2013) Memory, navigation and theta rhythm in the hippocampal-entorhinal system. Nat Neurosci 16:130–138.

Campbell EP, Martin L, Magee JC, Grienberger C (2026) Learning-dependent feedback by OLM interneurons shapes CA1 representations. bioRxiv.

Chadwick A, van Rossum MC, Nolan MF (2015) Independent theta phase coding accounts for CA1 population sequences and enables flexible remapping. Elife 4.

Deb K, Jain H (2013) An evolutionary many-objective optimization algorithm using reference-point-based nondominated sorting approach, part I: solving problems with box constraints. IEEE transactions on evolutionary computation 18:577–601.

Galloni AR, Peddada A, Chennawar Y, Milstein AD (2026) Cellular and subcellular specialization enables biology-constrained deep learning. Cell Rep 45:117159.

Geiller T, Vancura B, Terada S, Troullinou E, Chavlis S, Tsagkatakis G, Tsakalides P, Ocsai K, Poirazi P, Rozsa BJ, Losonczy A (2020) Large-Scale 3D Two-Photon Imaging of Molecularly Identified CA1 Interneuron Dynamics in Behaving Mice. Neuron 108:968–983 e969.

Geiller T, Sadeh S, Rolotti SV, Blockus H, Vancura B, Negrean A, Murray AJ, Rozsa B, Polleux F, Clopath C, Losonczy A (2022) Local circuit amplification of spatial selectivity in the hippocampus. Nature 601:105–109.

Geisler C, Diba K, Pastalkova E, Mizuseki K, Royer S, Buzsaki G (2010) Temporal delays among place cells determine the frequency of population theta oscillations in the hippocampus. Proc Natl Acad Sci U S A 107:7957–7962.

Gonzalez KC, Negrean A, Liao Z, Terada S, Zhang G, Lee S, Ocsai K, Rozsa BJ, Lin MZ, Polleux F, Losonczy A (2025) Synaptic basis of feature selectivity in hippocampal neurons. Nature 637:1152–1160.

Grienberger C, Magee JC (2022) Entorhinal cortex directs learning-related changes in CA1 representations. Nature 611:554–562.

Grienberger C, Chen X, Konnerth A (2014) NMDA receptor-dependent multidendrite Ca(2+) spikes required for hippocampal burst firing in vivo. Neuron 81:1274–1281.

Grienberger C, Milstein AD, Bittner KC, Romani S, Magee JC (2017a) Inhibitory suppression of heterogeneously tuned excitation enhances spatial coding in CA1 place cells. Nat Neurosci 20:417–426.

Grienberger C, Milstein AD, Bittner KC, Romani S, Magee JC (2017b) Inhibitory suppression of heterogeneously tuned excitation enhances spatial coding in CA1 place cells. Nat Neurosci.

Gritz S, Milstein AD (2026) Code repository for exploring spatial tuning of inhibition in CA1 place cells. https://github.com/Milstein-Lab/ca1_inh_tuning_models.

Hainmueller T, Cazala A, Huang LW, Bartos M (2024) Subfield-specific interneuron circuits govern the hippocampal response to novelty in male mice. Nat Commun 15:714.

Harnett MT, Makara JK, Spruston N, Kath WL, Magee JC (2012) Synaptic amplification by dendritic spines enhances input cooperativity. Nature 491:599–602.

Harvey CD, Collman F, Dombeck DA, Tank DW (2009) Intracellular dynamics of hippocampal place cells during virtual navigation. Nature 461:941–946.

Hunt MJ, Kasicki S (2013) A systematic review of the effects of NMDA receptor antagonists on oscillatory activity recorded in vivo. J Psychopharmacol 27:972–986.

Isaacson JS, Scanziani M (2011) How inhibition shapes cortical activity. Neuron 72:231–243.

Jaramillo J, Schmidt R, Kempter R (2014) Modeling inheritance of phase precession in the hippocampal formation. J Neurosci 34:7715–7731.

Jeong N, Singer AC (2022) Learning from inhibition: Functional roles of hippocampal CA1 inhibition in spatial learning and memory. Curr Opin Neurobiol 76:102604.

Lafourcade M, van der Goes MH, Vardalaki D, Brown NJ, Voigts J, Yun DH, Kim ME, Ku T, Harnett MT (2022) Differential dendritic integration of long-range inputs in association cortex via subcellular changes in synaptic AMPA-to-NMDA receptor ratio. Neuron 110:1532–1546 e1534.

Leung LS, Shen B (2004) Glutamatergic synaptic transmission participates in generating the hippocampal EEG. Hippocampus 14:510–525.

Li Y, Briguglio JJ, Romani S, Magee JC (2024) Mechanisms of memory-supporting neuronal dynamics in hippocampal area CA3. Cell 187:6804–6819 e6821.

Losonczy A, Magee JC (2006) Integrative properties of radial oblique dendrites in hippocampal CA1 pyramidal neurons. Neuron 50:291–307.

Marshall L, Henze DA, Hirase H, Leinekugel X, Dragoi G, Buzsaki G (2002) Hippocampal pyramidal cell-interneuron spike transmission is frequency dependent and responsible for place modulation of interneuron discharge. J Neurosci 22:RC197.

McKenzie S, Huszar R, English DF, Kim K, Christensen F, Yoon E, Buzsaki G (2021) Preexisting hippocampal network dynamics constrain optogenetically induced place fields. Neuron.

McNaughton BL, Barnes CA, O’Keefe J (1983) The contributions of position, direction, and velocity to single unit activity in the hippocampus of freely-moving rats. Exp Brain Res 52:41–49.

Milstein AD (2021) Code repository for nested: parallel multi-objective optimization software. https://github.com/neurosutras/nested.

Milstein AD, Tran S, Ng G, Soltesz I (2022) Offline memory replay in recurrent neuronal networks emerges from constraints on online dynamics. J Physiol.

Milstein AD, Li Y, Bittner KC, Grienberger C, Soltesz I, Magee JC, Romani S (2021) Bidirectional synaptic plasticity rapidly modifies hippocampal representations. Elife 10.

Mizuseki K, Sirota A, Pastalkova E, Buzsaki G (2009) Theta oscillations provide temporal windows for local circuit computation in the entorhinal-hippocampal loop. Neuron 64:267–280.

Nakazawa K, McHugh TJ, Wilson MA, Tonegawa S (2004) NMDA receptors, place cells and hippocampal spatial memory. Nat Rev Neurosci 5:361–372.

O’Keefe J, Conway DH (1978) Hippocampal place units in the freely moving rat: why they fire where they fire. Exp Brain Res 31:573–590.

O’Keefe J, Recce ML (1993) Phase relationship between hippocampal place units and the EEG theta rhythm. Hippocampus 3:317–330.

Okechukwu NG, Zaccone C, La Barbera L, Nobili A, D’Amelio M (2026) Excitation–inhibition imbalance as a common thread linking early Alzheimer’s disease with temporal lobe epilepsy. Experimental Neurology 397:115581.

Oppenheim AV, Schafer RW (2010) Discrete-time signal processing, 3rd Edition. Upper Saddle River: Pearson.

Poleg-Polsky A, Diamond JS (2016) NMDA Receptors Multiplicatively Scale Visual Signals and Enhance Directional Motion Discrimination in Retinal Ganglion Cells. Neuron 89:1277–1290.

Racca C, Stephenson FA, Streit P, Roberts JD, Somogyi P (2000) NMDA receptor content of synapses in stratum radiatum of the hippocampal CA1 area. J Neurosci 20:2512–2522.

Rolotti SV, Ahmed MS, Szoboszlay M, Geiller T, Negrean A, Blockus H, Gonzalez KC, Sparks FT, Solis Canales AS, Tuttman AL, Peterka DS, Zemelman BV, Polleux F, Losonczy A (2022) Local feedback inhibition tightly controls rapid formation of hippocampal place fields. Neuron 110:783–794 e786.

Shields BC et al. (2024) DART.2: bidirectional synaptic pharmacology with thousandfold cellular specificity. Nat Methods 21:1288–1297.

Skaggs WE, McNaughton BL, Wilson MA, Barnes CA (1996) Theta phase precession in hippocampal neuronal populations and the compression of temporal sequences. Hippocampus 6:149–172.

Spruston N, Jonas P, Sakmann B (1995) Dendritic glutamate receptor channels in rat hippocampal CA3 and CA1 pyramidal neurons. J Physiol 482 (Pt 2):325–352.

Valero M, Zutshi I, Yoon E, Buzsaki G (2022a) Probing subthreshold dynamics of hippocampal neurons by pulsed optogenetics. Science 375:570–574.

Valero M, Navas-Olive A, de la Prida LM, Buzsaki G (2022b) Inhibitory conductance controls place field dynamics in the hippocampus. Cell Rep 40:111232.

Virtanen P et al. (2020) SciPy 1.0: fundamental algorithms for scientific computing in Python. Nat Methods 17:261–272.

Weber SN, Sprekeler H (2018) Learning place cells, grid cells and invariances with excitatory and inhibitory plasticity. Elife 7.

Wilmes KA, Sprekeler H, Schreiber S (2016) Inhibition as a Binary Switch for Excitatory Plasticity in Pyramidal Neurons. PLoS Comput Biol 12:e1004768.

